# Stabilization of α-synuclein oligomers using formaldehyde

**DOI:** 10.1101/623538

**Authors:** Harm Ruesink, Lasse Reimer, Emil Gregersen, Arne Moeller, Cristine Betzer, Poul Henning Jensen

## Abstract

The group of neurodegenerative diseases, Parkinson’s disease (PD), dementia with Lewy bodies (DLB), and multiple system atrophy (MSA) all exhibit inclusions containing amyloid-type α-synuclein (α-syn) aggregates within degenerating brain cells. α-syn also exists as soluble oligomeric species that are hypothesized to represent intermediates between its native and aggregated states. These oligomers are present in brain extracts from patients suffering from synucleinopathies and hold great potential as biomarkers. Although easily prepared *in vitro*, oligomers are metastable and dissociate over time, thereby complicating α-syn oligomer research. Using the small amine-reactive cross-linker, formaldehyde (FA), we successfully stabilized α-syn oligomers without affecting their size, overall structure or antigenicity towards aggregate-conformation specific α-syn antibodies FILA and MJFR-14-6-4-2. Further, cross-linked α-syn oligomers show resistance towards denaturant like urea and SDS treatment and remain fully functional as internal standard in an aggregation-specific enzyme-linked immunosorbent assay (ELISA) despite prior incubation with urea. We propose that FA cross-linked α-syn oligomers could serve as important calibrators to facilitate comparative and standardized α-syn biomarker studies going forward.

## Introduction

Parkinson’s disease (PD), Dementia with Lewy Bodies (DLB) and Multiple System Atrophy (MSA) are the three main types of synucleinopathies, all of which are characterized by inclusions mainly consisting of intracellular aggregated α-synuclein (α-syn); a 140-amino acid protein with hypothesized unfolded native appearance [1, 2]. α-syn is widely expressed in the brain and mainly localizes to the presynaptic terminals where it is believed to play a role as a SNARE-complex chaperone, necessary for binding of vesicles and the release of neurotransmitter in the synaptic cleft [3, 4]. Proof of the involvement of α-syn in PD is evident from familial cases, where specific point mutations or multiple copies of the normal α-syn gene, SNCA, causes autosomal PD [5-8]. Under normal physiological conditions, α-syn exists as a native species in the brain, but aggregates into larger insoluble fibrils under pathological conditions [9]. However, α-syn also exists in an intermediate state, as an aggregated soluble species, called oligomers. Oligomers have been found in post-mortem brain extracts from both DLB, PD and MSA patients [10-12], and accumulating evidence indicates that oligomers are responsible for the toxicity seen in the brain of patients suffering from synucleinopathies [13-15]. Furthermore, *in vitro* generated oligomers are capable of seeding and mediate cell-to-cell spreading linked to the prion-like spreading seen in PD [16]. α-syn oligomers can be generated in vitro in several ways, such as chemical modification by dopamine, ethanol and Fe^3+^ and high protein concentration [17-19]. Oligomers formed spontaneously at high protein concentration have also shown similarities with patient-derived *in vivo* oligomers in the antigenicity towards aggregation-specific antibodies [10]. However, these oligomers are metastable and dissociate into monomers over time [20]. Therefore, an *in vivo* representative α-syn oligomer-standard used to study the underlying mechanistic pathways and aid in the search for synucleinopathy-specific biomarkers, is of great interest. Using formaldehyde (FA) crosslinking, we have successfully stabilized α-syn in its oligomeric state without disturbing antigenicity and biochemical properties, thereby providing a much-needed calibrating tool, which enables comparative and standardized research within the field of oligomeric α-syn.

## Results

### Characterizing native α-syn oligomers

Brief incubating of high concentration of α-syn at 0°C has previously been shown to spontaneously generate oligomeric species [19-21]. Using this method, in combination with size-exclusion chromatography, we successfully separated oligomers from monomers and obtained a pure oligomeric fraction (Fig 1A). Dynamic light scattering (DLS) confirmed the different sizes of the α-syn monomers and oligomers, showing two distinct species, with an average radius of 3.6 ± 0.3 and 49.7 ± 2.6 nm respectively (Fig. 1B). Monomers and oligomers both bind the α-syn specific antibodies BD and ASY1 that recognizes total α-syn levels, whereas only the oligomers specifically binds the two aggregation-specific antibodies, FILA and MJF14 (Fig. 1C). Transmission electron microscopy (TEM) revealed a twisted ribbon-like structure of the purified α-syn oligomers, which corresponds well with previous observation of oligomeric α-syn (Fig. 1D) [22]. The α-syn oligomers clearly differ from the well-ordered structure and size of the preformed α-syn fibrils (Fig. 1E).

**Figure 1.**
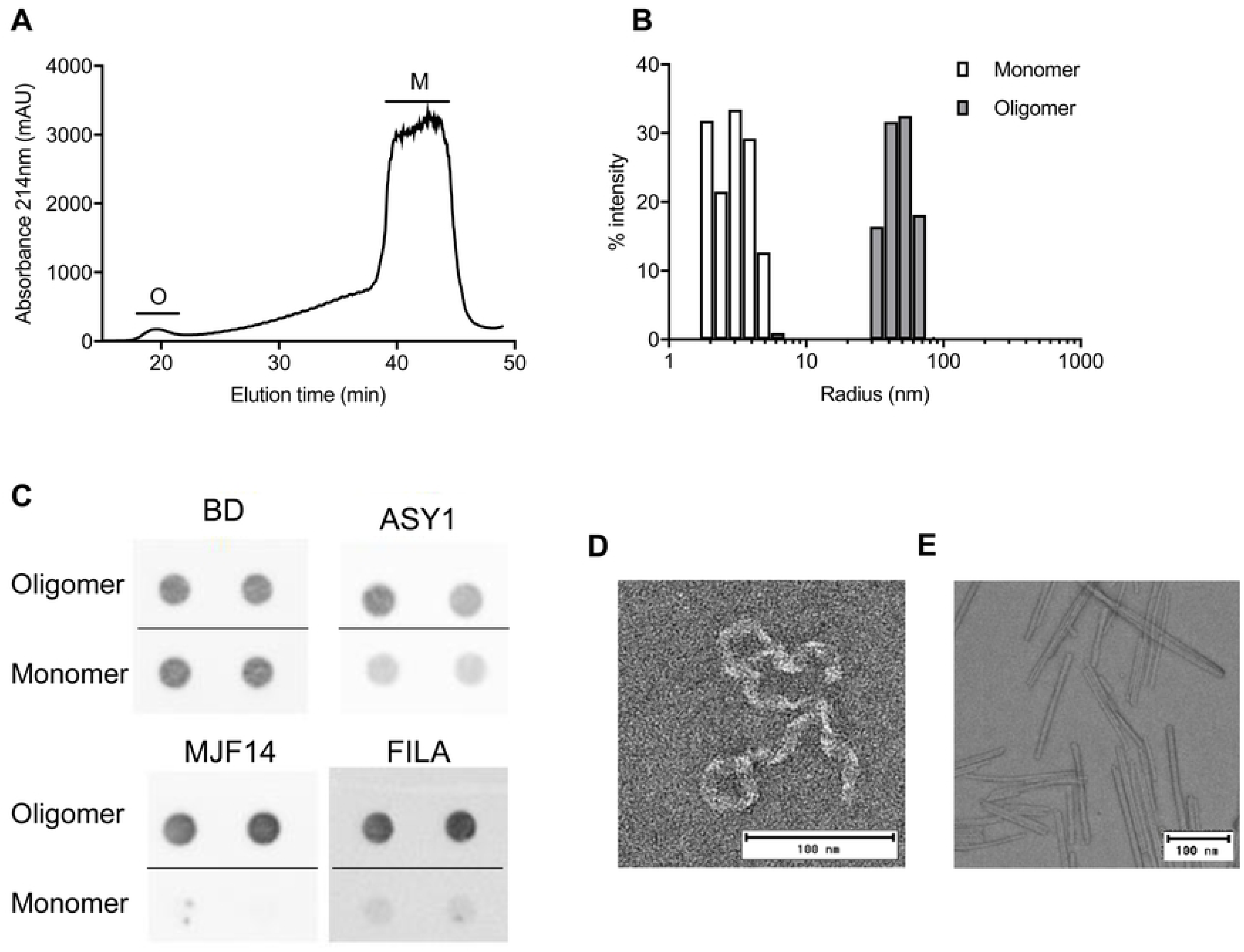
Generation and confirmation of in vitro generated α-syn oligomers and fibrils. **A)** Oligomers generated by resuspending lyophilised recombinant α-syn at high concentration (10mg/mL) incubated on ice and subsequently isolated using gel-filtration. Oligomers (O) are collected between 18-22 min. and monomers (M) between 38-43 min. **B)** Hydrodynamic radius in nm of isolated particles was determined using DLS. Graph shows a merged image of the hydrodynamic radius (x-axis) of oligomers (dark grey) and monomers (white). Intensity of signal is depicted on the y-axis **C)** Antigenicity of oligomers and monomers to BD, FILA5, MJF14 and ASY1 was determined using dot blot. Dots consist of 50ng protein spotted in duplicates. Negative stain EM image shows ultrastructure of native α-syn oligomer **D)** and **E)** ultrastructure of in vitro formed α-syn fibrils.

### Stabilization of the α-syn oligomers using formaldehyde

α-syn oligomers consist of non-covalently bound monomers that easily dissociate into monomers upon boiling in SDS prior to SDS-PAGE (Fig. 2A, right). However, incubation of the α-syn oligomers with increasing concentration of the small amine-reactive cross-linker, formaldehyde (FA), prior to SDS-PAGE, stabilized the α-syn multimers as evident from a depletion of the ∼17kDa monomeric α-syn band (Fig. 2A, right). The retention of immunoreactivity in the stacking gel demonstrates the cross-linking of α-syn into large stable complexes (Fig 2A, right). By contrast α-syn monomers incubated with identical FA concentrations did not result in cross-linked-dependent depletion of the monomeric band nor accumulation of high molecular weight species (Fig 2A, left).

**Figure 2.**
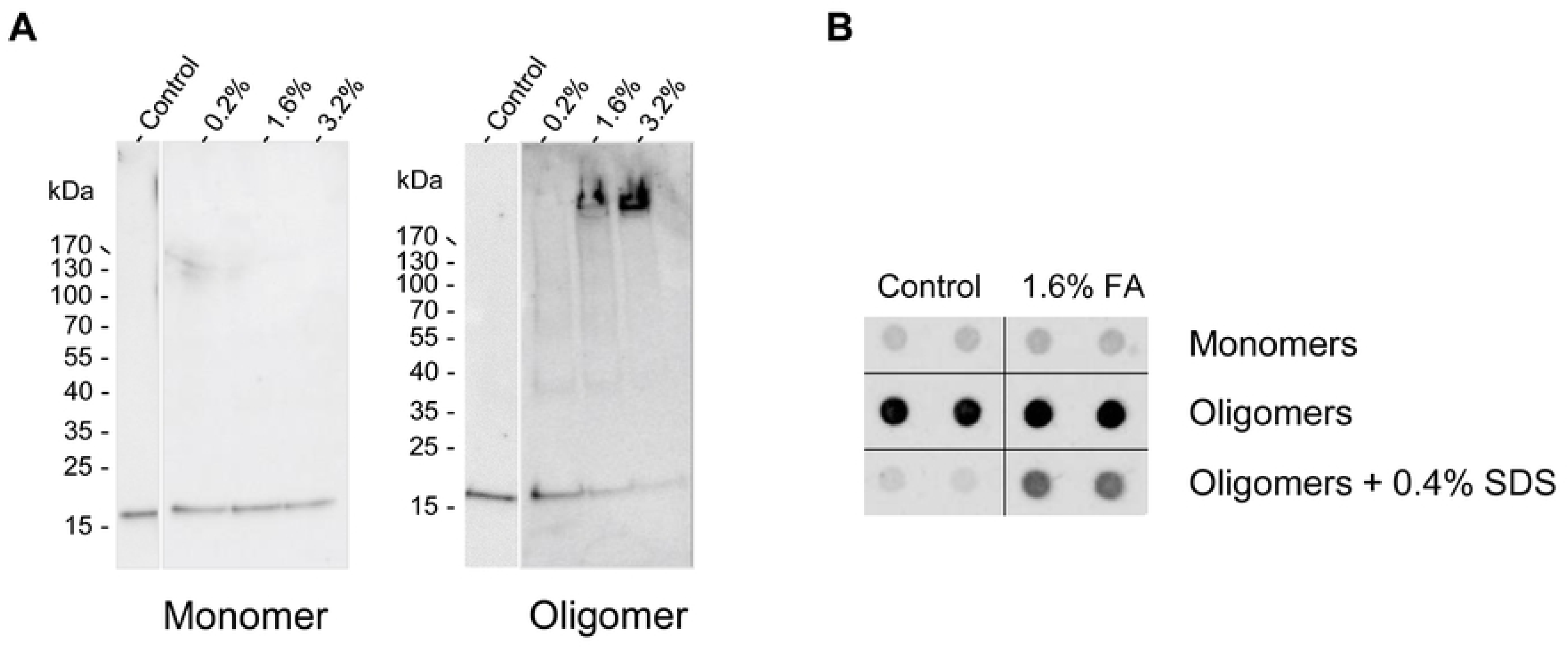
Crosslinking of α-syn monomers and oligomers. **A)** α-syn monomers and oligomers were crosslinked with FA at different concentrations. Immunoblot of monomers (left) and oligomers (right) show ASY1 binding. Monomeric α-syn situates at ∼17kDa. Depletion of the ∼17kDa α-syn band and presence of ASY-1 reactivity in the stacking gel suggest efficient cross-linking upon FA treatment of oligomers. **B)** Dot blot of 100ng non-treated- or 1.6% FA cross-linked α-syn monomers and oligomers using aggregation-specific FILA5 antibody. Prior to dot-blot, one subset of native- and cross-linked oligomers were treated with 0.4% SDS for 1 h. to assess oligomer stability.

Visualization of FA-treated monomeric and oligomeric α-syn, using the aggregation-specific FILA antibody, revealed that epitope-recognition was unaffected by cross-linking (Fig. 2B). The FA treated oligomers are also less prone to disassembly, as FILA antigenicity after 3h incubation with denaturizing SDS, is still observed (Fig. 2B).

### Optimization of cross-linking

To avoid excessive and unnecessary chemical modification of the oligomers, we carried out titration experiments, aiming to find the lowest possible concentration of FA that efficiently stabilize α-syn oligomers. Using 0.2-3.2% FA and incubation for 30- or 60 min, we found that 1.6% and 3.2% FA after 60 min resulted in satisfactory cross-linking as measured by loss of the α-syn monomer-band (95 ± 4.6% and 98 ± 1.9% respectively), when resolved on SDS-PAGE (Fig. 3A-C). Further, we assessed if antigenicity of α-syn monomers or oligomers were affected by FA cross-linking. No difference was observed in oligomer recognition using antibodies BD, ASY1, FILA and MJF14 at any concentration of FA (Fig. 3D-G), although a trend of high FA concentrations masking the BD epitope of monomeric α-syn was observed (Fig. 3E). 3.2% FA incubation generally results in a slightly higher cross-linking efficiency than 1.6% FA (Fig. 3A and 3C), but we chose to continue with 1.6% FA for the remainder of the study for technical reasons. Primarily to avoid unnecessary dilution of the oligomers by adding larger volumes of Tris to quench the FA cross-linker.

**Figure 3.**
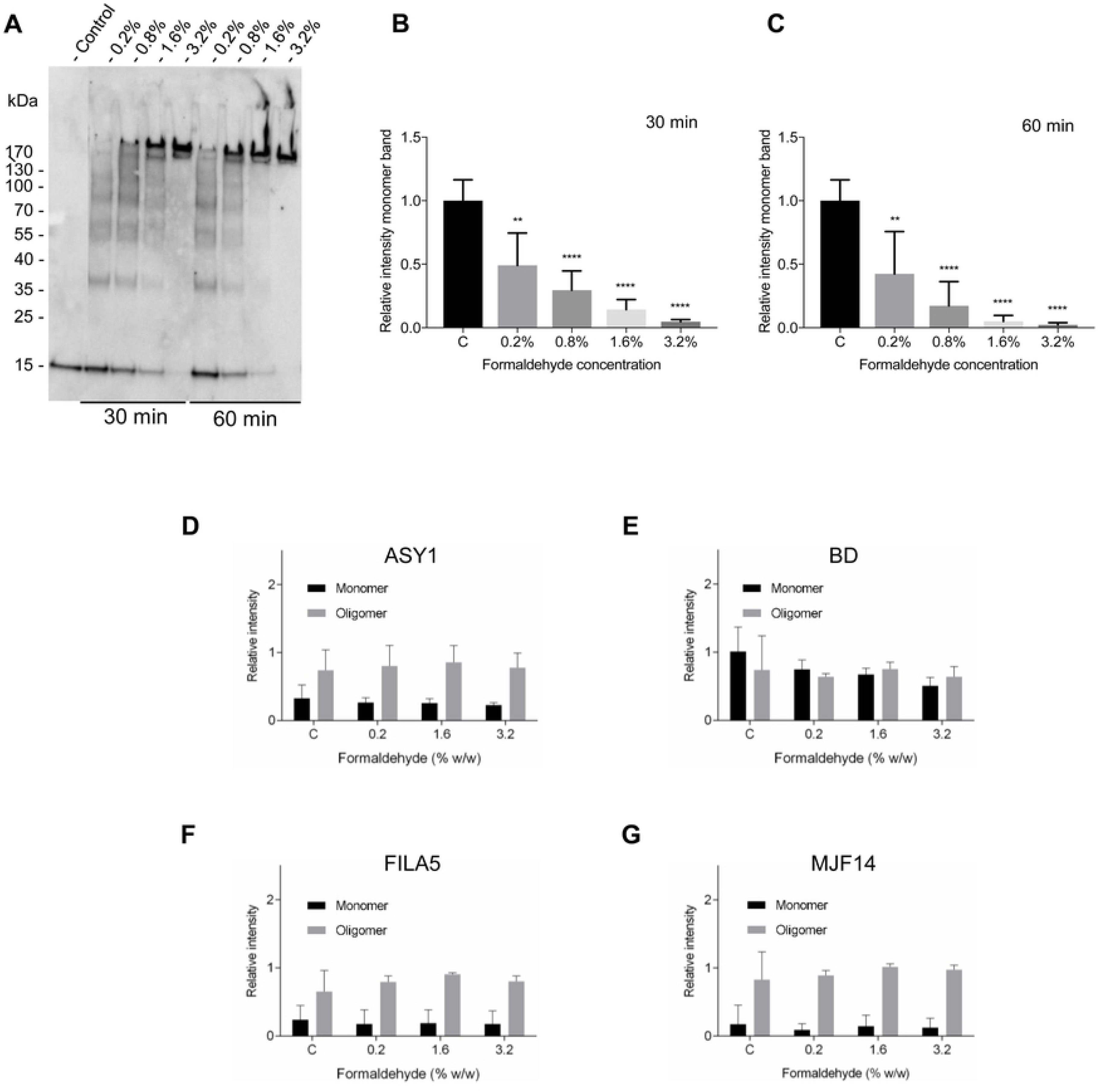
Optimization of α-syn oligomer cross-linking. **A-C)** α-syn oligomers were crosslinked with FA in a concentration of 0-3.2% for 30- or 60 min and subjected to immunoblot using ASY1 antibody to detect α-syn species (A). Monomer ∼17kDa ASY-1 positive bands were quantified to assess degree of cross-linking and results are shown in B and C. Bar graphs show means ± SD obtained in three individual experiments. **D-G)** Monomers (black bars) and oligomers (grey bars) were cross-linked using different FA concentrations (0%-3.2% FA) for 60 min. The antigenicity of cross-linked monomers and oligomers were assessed via dot blot using ASY1 (D), BD (E), FILA5 (F) and MJF14 (G) antibodies. Figures show means ± SD of three independent experiments. *p<0.05, **p<0.01, ***p<0.001, ****p<0.0001, n.s.= not significant.

### PFA stabilized α-syn retain biophysical and experimental properties

To test the biophysical properties of the 1.6% FA cross-linked oligomers, we performed a DLS time-course experiment, measuring the size-distribution of the oligomers after 0, 15, 30 and 60 min of 1.6% FA treatment. Throughout the experiment, the size of the oligomers was unchanged, suggesting that the oligomers are stabilized in their native structure and not cross-linked into larger complexes over time (Fig. 4A). EM confirmed that the overall structure of the oligomers were kept after FA mediated cross-linking as these resembled native oligomers (Fig. 1D and 4B).

**Figure 4.**
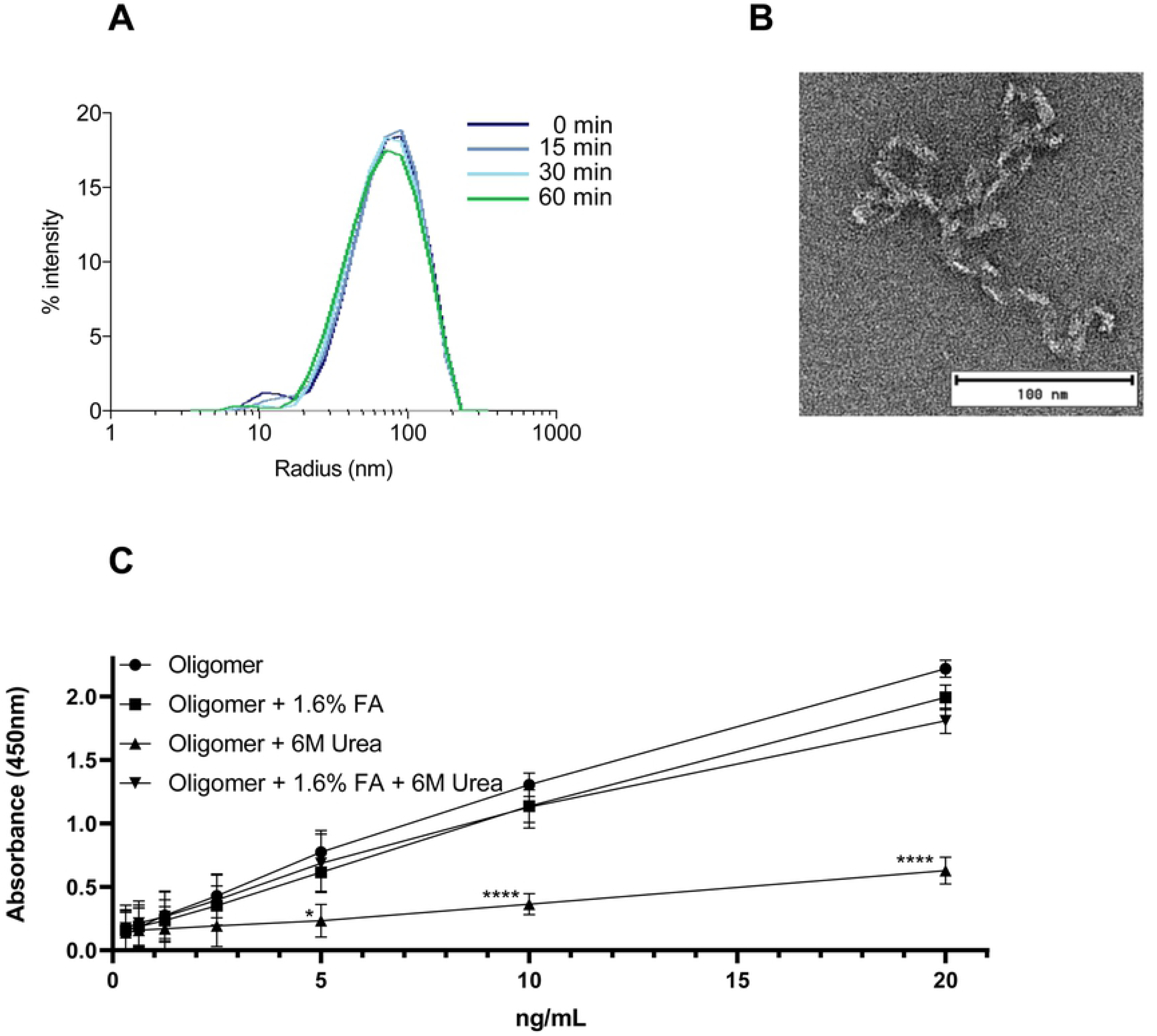
Oligomers cross-linked using FA maintain their size and structure and resist denaturation. **A)** Hydrodynamic radius of oligomers was monitored during 1.6% FA cross-linking using DLS. Measurements were taken before FA treatment (0 min) and 15-, 30- and 60 min after initiation of cross-linking. **B)** Negative stain EM image shows ultrastructure of α-syn oligomer cross-linked for 60 min using 1.6% FA. **C)** Native and 1.6% FA crosslinked α-syn oligomer were treated with 6M Urea for 6h at RT or left untreated. Serial dilutions were analyzed by ELISA as previously described by Lassen et al. [23], utilizing the aggregated specific antibody MJF14. The levels are measured as absorbance at 450 nm. Two-way ANOVA for repeated measures followed by Tukey’s multiple comparison test was used for ELISA experiment.

*In vitro* formed α-syn oligomers are useful in many experimental setups. One example, is the application of *in vitro* α-syn oligomers as internal standard in the newly developed aggregation-specific enzyme-linked immunosorbent assay (ELISA) based on the aggregation-specific MJF14 antibody [23]; an assay that effectively assess levels of α-syn oligomers in cell and animal models. Using this method, we compared the standard curves prepared from either native oligomers or FA cross-linked oligomers under different conditions (Fig. 4C). We observed that ELISA standard curves using cross-linked oligomers were indistinguishable from those prepared with native oligomers (Fig. 4C). Moreover, while pretreatment of native oligomers with 6M urea clearly abolish the MJF14 ELISA signal, FA cross-linked oligomers remained stable and created reliable standard curves despite exposure to high concentration urea (Fig. 4C). Together, these findings demonstrate that FA treatment effectively cross-link and stabilize native oligomers without disturbing size, structure and antibody recognition of the oligomers.

## Discussion

Mounting evidence suggest a central role of soluble α-syn oligomers as toxic species relevant to the progressive spreading of α-syn pathology in the nervous system in synucleinopathies [24] and quantification of such species holds great potential as biomarkers [25-27].

In vitro formed α-syn oligomers represents a very heterogenous group that either presents as annular-, spherical-, circular-, rod shaped-, amorphous- and twisted ribbon-like structures dependent on the method of preparation and visualization [19, 21, 28]. Their diverse size, biochemical and structural properties likely make them dissimilar with respect to their presentation of antigens towards conformation specific-antibodies as demonstrated for insoluble α-syn strains [28, 29]. Moreover their semistable nature makes them unsuitable as analytical calibration standards in bioassays [20].

We successfully cross-linked α-syn oligomers using FA, thereby increasing stability towards SDS and urea denaturation. This oligomer preparation binds neuronal proteins in a conformational specific manner [21, 22], and also bind the two conformational specific antibodies FILA and MJF14. The FA mediated crosslinking did not disturb the overall twisted ribbon-like oligomeric structure, the size of the oligomers, their antigenicity toward FILA and MJF14 or the crosslinked oligomers applicability as calibrators in an α-syn oligomer specific ELISA assay [23]. Hence we propose this FA cross-linked α-syn oligomer preparation could serve as calibrators that facilitates comparative and standardized biomarker studies in the synucleinopathies.

## Materials and methods

### Antibodies

Primary antibodies: α-syn monoclonal mouse-anti-α-syn (BD Biosciences, catalog no. 610787), polyclonal rabbit-anti-α-syn (ASY1 [30]), α-syn aggregate-specific antibodies FILA5 (polyclonal rabbit-antibody, in house generated [31]), and MJFR-14-6-4-2 (MJF14) (abcam, catalog no. 209538, rabbit monoclonal antibody, generated against full length α-syn protein filament). Secondary antibodies: horseradish peroxidase (HRP)-conjugated polyclonal rabbit-anti-mouse (Dako, P0260), HRP-conjugated polyclonal swine-anti-rabbit (Dako, P0217).

### Oligomer preparation

Production of α-syn and α-syn oligomers was performed as previously described [19, 21, 32]. In short, purified α-syn was dissolved in PBS to a final concentration of 10mg/mL and incubated on ice for 30 minutes while vortexed regularly. After incubation, α-syn was centrifuged at 12000g for 5 minutes and the supernatant loaded onto a Superdex™ 200 10/300 GL column (GE Healthcare). The column was eluted with PBS at a flow rate of 0.5 mL/minute. Oligomers were collected between 18-22 min and monomers after 38-43 min. All collected fractions were snap frozen on dry ice and stored at −80°C.

### Crosslinking

Monomers and oligomers were crosslinked at a final protein concentration of 10-15ng/μL. Formaldehyde 37-38% w/w (PanReac AppliChem) was diluted in H_2_O and filtered (0,45μm) before use. FA crosslinker was added in a 2x concentration to samples to obtain final concentration. Crosslinking was performed at 37°C and quenched with Tris (two times molar concentration of FA) for a minimum of 15 minutes at RT prior to dialysis in PBS.

### Immunoblotting

For Western blotting, samples were boiled for 5 min at 95°C for in loading buffer (50 mM Tris pH 6.8, 40% glycerol, 4% SDS, bromophenol blue) without reducing agent before loading onto a 4-16% PAGE gel (GenScript ExpressPlus™). Gels were blotted on PVDF membranes and fixed in 4% formaldehyde for 30 minutes and boiled in PBS for 5 minutes. For dot blotting 50-100ng of protein was blotted on nitrocellulose (Hydrobond-C Extra, GE Healthcare) and these filters were not fixed or boiled. All membranes were blocked in non-fat milk (TBS with 5% non-fat milk, 0.05% Tween 20 and 0.02% sodium azide) for 30 minutes at RT followed by incubation with primary antibodies overnight. membranes were washed in TBS-T (TBS with 0.1% Tween) and incubated with secondary antibodies for 1.5 hour. Membranes were washed, developed using ECL reagent (Pierce™, Thermo Scientific), and subsequently imaged on a LAS-3000 imaging system (Fuji). Blot quantification was done using ImageJ

### Size determination using Dynamic Light Scatter

For size determination using Dynamic Light Scattering (DLS) samples were placed in a disposable cuvette and analyzed using a Wyatt Technology DynaPro NanoStar. Laser strength was set to 100% and scatter angle fixed at 90°. Total samples measurement where assembled via 10 consecutive 5 second measurements. Prior to analysis, samples were centrifuged for 3 minutes at >12000g. Data was analyzed with Dynamics V7.5.0.17.

### Negative staining transmission electron microscopy

Negative staining transmission electron microscopy (TEM) was performed as described previously [33, 34]. In brief, 3μL of sample was loaded onto a glow discharged carbon coated copper grid and stained with 2% uranyl formate. Images were collected using a Tecnai G2 Spirit TWIN electron microscope in combination with a Tietz TemCam-F416 CMOS camera at a magnification of 54000x to 67000x. Samples for EM were crosslinked in PBS and dialyzed, with three buffer changes, to TBS to avoid precipitation of phosphate with uranyl. Sample to dialyzing buffer volume ratio 1:100.

### Enzyme-linked immunosorbent assay

ELISA analysis was carried out as previously described [23]. In summary, 62.5 ng/mL MJF14 antibody and 0.5 µg/mL BD antibody were used as capture and detection antibody, respectively. 1,1 µg/mL native and crosslinked α-syn oligomer were incubated without or in the presence of 6 M urea for 6 hours at RT, before being analyzed by ELISA in at dilution ranging from 20 ng/mL to 313 pg/mL.

### Statistical analysis

Data was statistically analyzed and visualized using GraphPad Prism 7. Data was tested with a one-way ANOVA followed by Bonferroni post hoc analysis. For ELISA experiment, two-way ANOVA for repeated measures followed by Tukey’s multiple comparison test was used. Differences were considered significant for *p<0.05, **p<0.01, ***p<0.001, ****p<0.0001. Data is presented as means ± SD.

## Acknowledgement

The study was supported by Michael J Fox grant 13750 and Lundbeck foundation grants R223-2015-4222, R248-2016-25, R248-2016-2518 Danish Research Institute of Translational Neuroscience - DANDRITE, Nordic-EMBL Partnership for Molecular Medicine

